# Discovery of molecular features underlying morphological landscape by integrating spatial transcriptomic data with deep features of tissue image

**DOI:** 10.1101/2020.06.15.150698

**Authors:** Sungwoo Bae, Hongyoon Choi, Dong Soo Lee

**Author notes:** **Correspondence** Hongyoon Choi, MD. Ph.D. Department of Nuclear Medicine, Seoul National University Hospital, 28 Yongon-Dong, Jongno-Gu, Seoul, 110-744, Korea, Dong Soo Lee, MD., Ph.D. Department of Nuclear Medicine, Seoul National University Hospital, 28 Yongon-Dong, Jongno-Gu, Seoul, 110-744, Korea.

## Abstract

Profiling molecular features associated with the morphological landscape of tissue is crucial to interrogate structural and spatial patterns that underlie biological function of tissues. Here, we present a new method, SPADE, to identify important genes associated with morphological contexts by combining spatial transcriptomic data with co-registered images. SPADE incorporates deep learning-derived image patterns with spatially resolved gene expression data to extract morphological context markers. Morphological features that correspond to spatial maps of transcriptome were extracted by image patches surrounding each spot, and then, represented by image latent features. The molecular profiles correlated with the image latent features were identified. The extracted genes could be further analyzed to discover functional terms and also exploited to extract clusters maintaining morphological contexts. We apply our approach to spatial transcriptomic data of three different tissues to suggest an unbiased method capable of offering image-integrated gene expression trends.

## Main

Until recently, multitudes of technologies have been developed to analyze spatial gene expression patterns that provide molecular profiling with spatial information in tissues^1^. In particular, recent progress in spatial gene expression technologies which applies next-generation sequencing with spatial barcode, fluorescence in situ hybridization (FISH), or in situ sequencing (ISS) have innovated experimental approaches to decipher the spatial heterogeneity of biological process^2-5^. A spatial context at single-cell level resolution has allowed the analysis of the location of heterogeneous cells and their spatial interactions in tumor tissues as well as brain, the human heart, and inflammatory tissues^4, 6-11^.

Even though spatial gene expression analyses have been actively developed and applied to various tissues and diseases, analytic methods that integrate transcriptome and imaging data are lacking. In spite of the feasibility of analysis that combines gene expression, spatial interaction between different spots of spatial barcodes and image patterns, most methods have regarded gene expression from spots as independent samples and interpreted like single-cell RNA-sequencing (scRNA-seq) data^4, 6-9^. Especially, one of the advantages of spatial gene expression data is additional information of co-registered images which contains morphological as well as functional patterns. In this regard, important genes related to the image features can be extracted and further utilized to interrogate molecular profiles underlying structural and morphological architectures.

Here we introduce a method for identifying **spa**tial gene expression patterns by **de**ep learning of tissue images (**SPADE**). SPADE extracts gene expression markers by incorporating morphological patterns of an image patch surrounding each spot which contains transcriptomic data. A convolutional neural network (CNN) was employed to define image latent features associated with gene expression. We present molecular markers of various tissues associated with the morphological landscape to discover not only a spatial trend of gene expression in tissues but also biological processes related to histologic architecture.

## Results

### Markers associated with morphological landscape for breast cancer tissue

We discovered important gene markers correlated to image features extracted by a CNN (**Fig. 1a**). Three publicly available dataset were analyzed to identify gene expression markers associated with morphological landscape, defined as SPADE genes. Image latent features represented by 512-dimensional vectors were extracted by a pretrained CNN model, VGG-16^12^, from image patches surrounding spots which correspond to transcriptome data. To define highly variable image latent features, principal component analysis (PCA) was applied to the output of the VGG-16 for all patches corresponding to spots. SPADE genes were identified among highly variable genes (HVG) by linear regression analysis with principal components of image features. We also used SPADE genes for further analyses including functional gene ontology to characterize biology in tissues and clustering of spots.

**Fig. 1:**
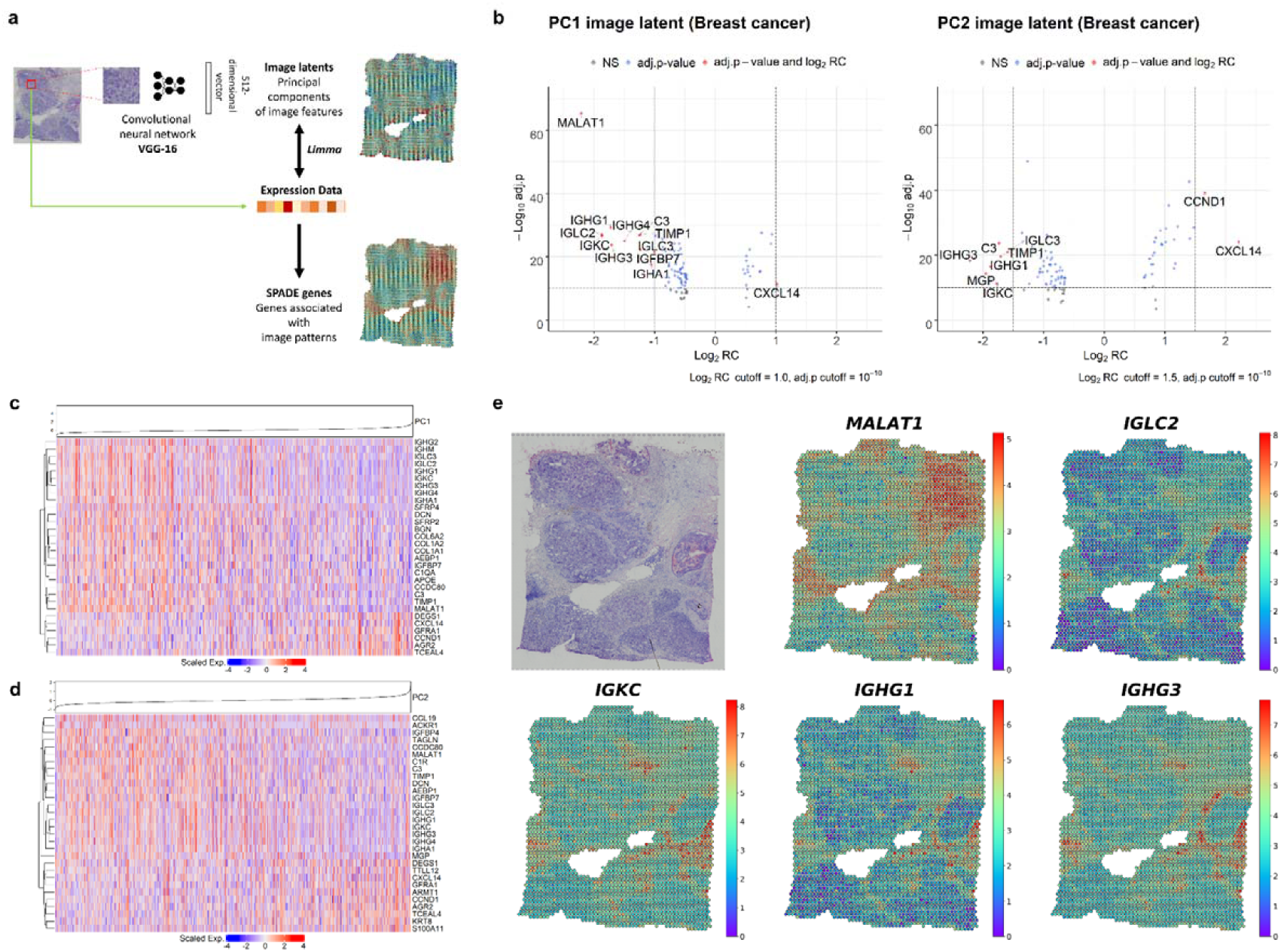
Discovery of image-integrated spatially variable genes in breast cancer data. a, Multiple patches were extracted from a tissue slide image based on the coordinates of sampling spots. Each image patch was provided as an input to pretrained convolutional neural network (CNN) model, VGG-16. 512 image features extracted from the CNN were further processed with principal component analysis (PCA) to reduce the dimensions. SPADE genes were constructed by a linear model to identify gene expression correlated with PC image latent of each spot. b, Volcano plots for highly associated genes with PC1 and PC2 image latent features. Cutoff for log_2_ regression coefficient (RC) is 1.0 in PC1 and 1.5 in PC2. Cutoff for adjusted p-value (Benjamini-Hochberg correction) is 10^−10^ for both PC1 and PC2. c, A heatmap for top 30 highly associated genes for log_2_RC in PC1 image latent space were drawn. Hierarchical clustering was performed for top 30 genes and PC1 value in each of the spot was presented on top. d, A heatmap for top 30 highly associated genes for log_2_RC in PC2 image latent space. Hierarchical clustering was performed for top 30 genes and PC2 value in each of the spot was shown on top. e, Spatial expression of top 5 genes representing greatest contrast in PC1 image latent space. Gene expression level of each spot was visualized with a colormap.

Firstly, SPADE was applied to human breast cancer tissue containing 3813 sampling spots. Dimension of 512 image features were reduced by PCA and principal component 1 and 2 (PC1 and PC2) values of each spot were mapped on H&E slide (**Supplementary Fig. 1**). Genes associated with PC1 and PC2 were identified (**Fig. 1b, Supplementary Fig. 2**). Top 30 genes with false discovery rate (FDR) below 0.05 were selected and then represented as a heatmap according to the increase of PCs (**Fig. 1c, d)**. The top 5 genes *MALAT1, IGLC2, IGKC, IGHG1*, and *IGHG3* from PC1 image latent were mapped on the tissue (**Fig. 1e**). Top 5 genes from PC2 image latent also showed the spatially variable expression according to image features (**Supplementary Fig. 3**).

### Functional molecular features associated with image-based landscape of breast cancer tissue

Gene ontology (GO) analysis^13, 14^ was performed with SPADE genes derived from PCs. From PC1 to PC6 explained 50.45%, 22.93%, 8.47%, 5.56%, 3.54%, and 1.53% of data variance in 512-dimensional image latent, respectively. PC1, PC2, and PC5 SPADE gene sets were enriched with extracellular matrix and various immune response GO terms in molecular function, cellular component, and biological process subcategories. On the other hand, PC3 SPADE genes over-represented GO terms regarding response to metal ion and PC6 over-represented biological response to hypoxia (**Fig. 2a, Supplementary Fig. 4**).

**Fig. 2:**
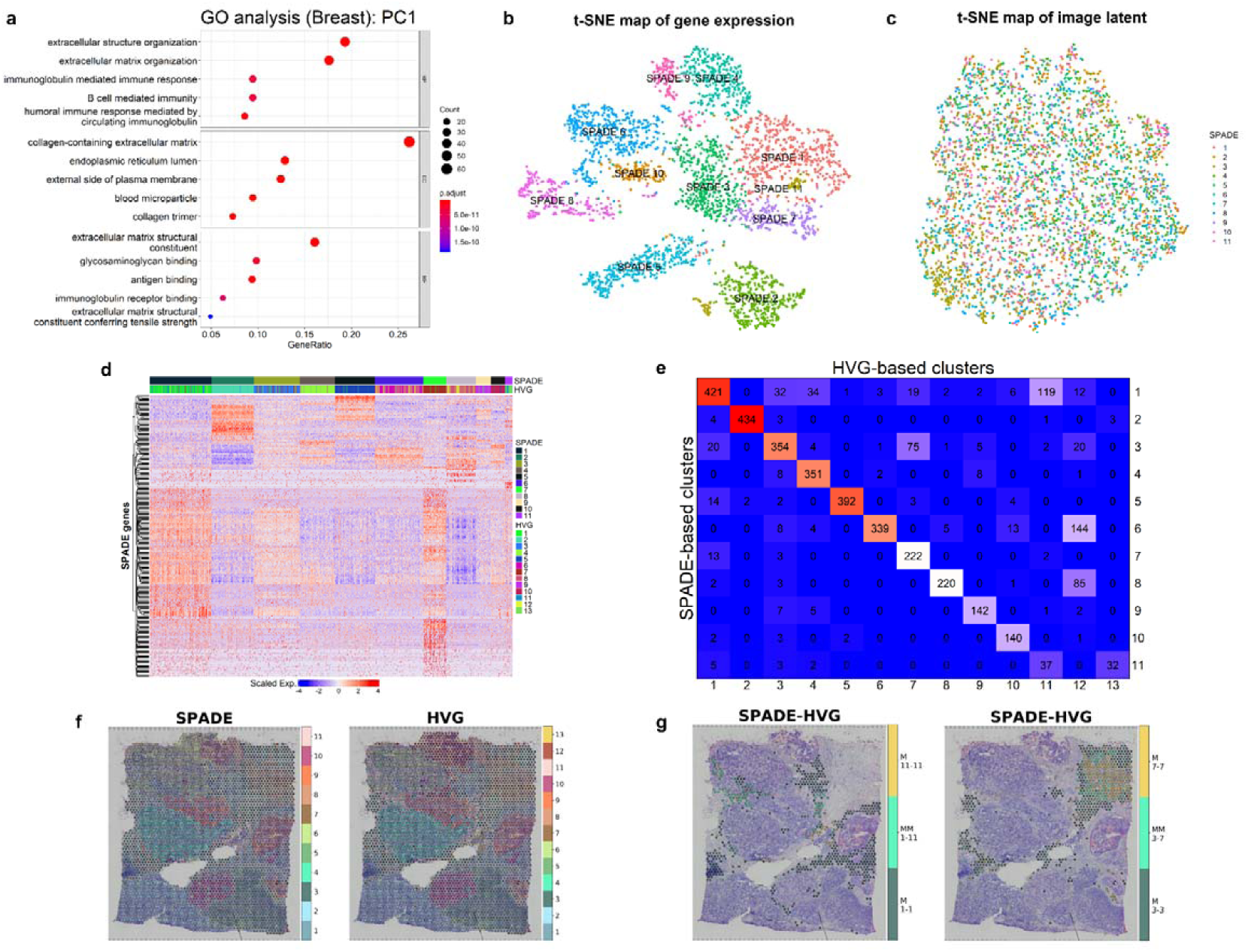
Functional molecular features of SPADE and spot clustering in breast cancer data. a, Gene ontology (GO) analysis for SPADE genes from PC1 image latent showing top 10 over-represented gene functions in each subcategory, molecular function (MF), cellular component (CC), and biological process (BP). Number of overlapped genes was expressed as size of dot and Benjamini-Hochberg adjusted p-value was exhibited with colormap. b, t-SNE plot of transcriptomic data. SPADE genes were utilized to classify spots into 11 SPADE-based clusters. c, t-SNE plot for deep learning-derived image features. SPADE-based cluster identity of each spot was visualized. d, A heatmap for expression of SPADE genes in every sampling spot. SPADE and HVG-based cluster identity of each spot was annotated on top of heatmap. e, A cross table exhibiting number of overlapped spots in corresponding SPADE and HVG-based clusters. f, Spatial distribution of clusters based on SPADE genes or HVG mapped on the tissue slide. g, Spatial mapping of the matched and mismatched spot clusters. Results for SPADE 1-HVG 11 is on the left and SPADE 3-HVG 7 is on the right. M is abbreviation for matched cluster and MM is for mismatched cluster.

### Clustering based on SPADE genes in breast cancer tissue

For the next step, spots were clustered using SPADE genes. 350 SPADE genes were selected from PC1 and PC2 image latent and spots were clustered with shared nearest neighborhood (SNN) graph^15, 16^. The clusters based on SPADE genes were visualized by two-dimensional t-distributed stochastic neighbor embedding (t-SNE)^17^ of transcriptomic data **(Fig. 2b)** and image latent (**Fig 2c**). The markers of clusters were identified (**Supplementary Fig. 5a**). As conventional methods use HVG instead of image-related genes, the patterns of clusters were similar but different. HVG-based clusters were represented by t-SNE map and also markers for each cluster were extracted (**Supplementary Fig. 5b, c)**. A heatmap for the SPADE genes revealed distinct gene expression patterns in each SPADE-based and HVG-based cluster (**Fig. 2d**). The percentage of shared spots of SPADE-based with HVG-based clusters were ranged from 46.84% (cluster 11: 37/79) to 97.53% (cluster 2: 434/445) (**Fig. 2e**). The number of HVG, SPADE genes, and marker genes of clusters derived by HVG and SPADE were represented as a Venn-diagram (**Supplementary Fig. 5d**). The spatial mapping of the SPADE and HVG-based clusters were exhibited (**Fig. 2f**). In addition, the top 2 mismatched clusters based on SPADE and HVG with the highest number of spots were represented. The spots classified as SPADE 1-HVG 11 cluster and SPADE 3-HVG 7 cluster were mapped on the tissue slide (**Fig. 2g**). When the spots were represented by PCs of image latent, spots with mismatched clusters were closer to the SPADE-based cluster according to the distance measured on image latent (**Supplementary Fig. 6**). The distance from mismatched spots (SPADE 1-HVG 11) to SPADE-based cluster (median distance: 0.38) was significantly shorter than the distance to HVG-based cluster (median distance: 0.66) (p<0.001, Mann-Whitney U test). Another mismatched cluster, SPADE 3-HVG 7, was more closely located to SPADE-based cluster (median distance: 0.32) than the HVG-based cluster (median distance: 0.36), although it was not significant (p=0.108).

### Markers related to morphological patterns of olfactory bulb and kidney tissues

Utility of SPADE was further validated with mouse olfactory bulb tissue analyzed by a spatial transcriptomics paper^4^ and mouse kidney tissue from 10x genomics. Mouse olfactory bulb included 267 spots and mouse kidney data contained 1438 spots with spatial gene expression profiles. PC1 and PC2 values in each spot were mapped on the tissue (**Supplementary Fig. 7)**. Results for linear regression analysis were presented as volcano plot (**Fig. 3a, b)** and scatter plot (**Supplementary Fig. 8**). In addition, spatial marker genes showing greatest regression coefficient in PCs were identified. Top 30 genes for PC1 and PC2 image latent in olfactory bulb and kidney exhibited marked contrast of gene expression as the PC1 and PC2 value changes (**Fig. 3c, d, Supplementary Fig. 9**).

**Fig. 3:**
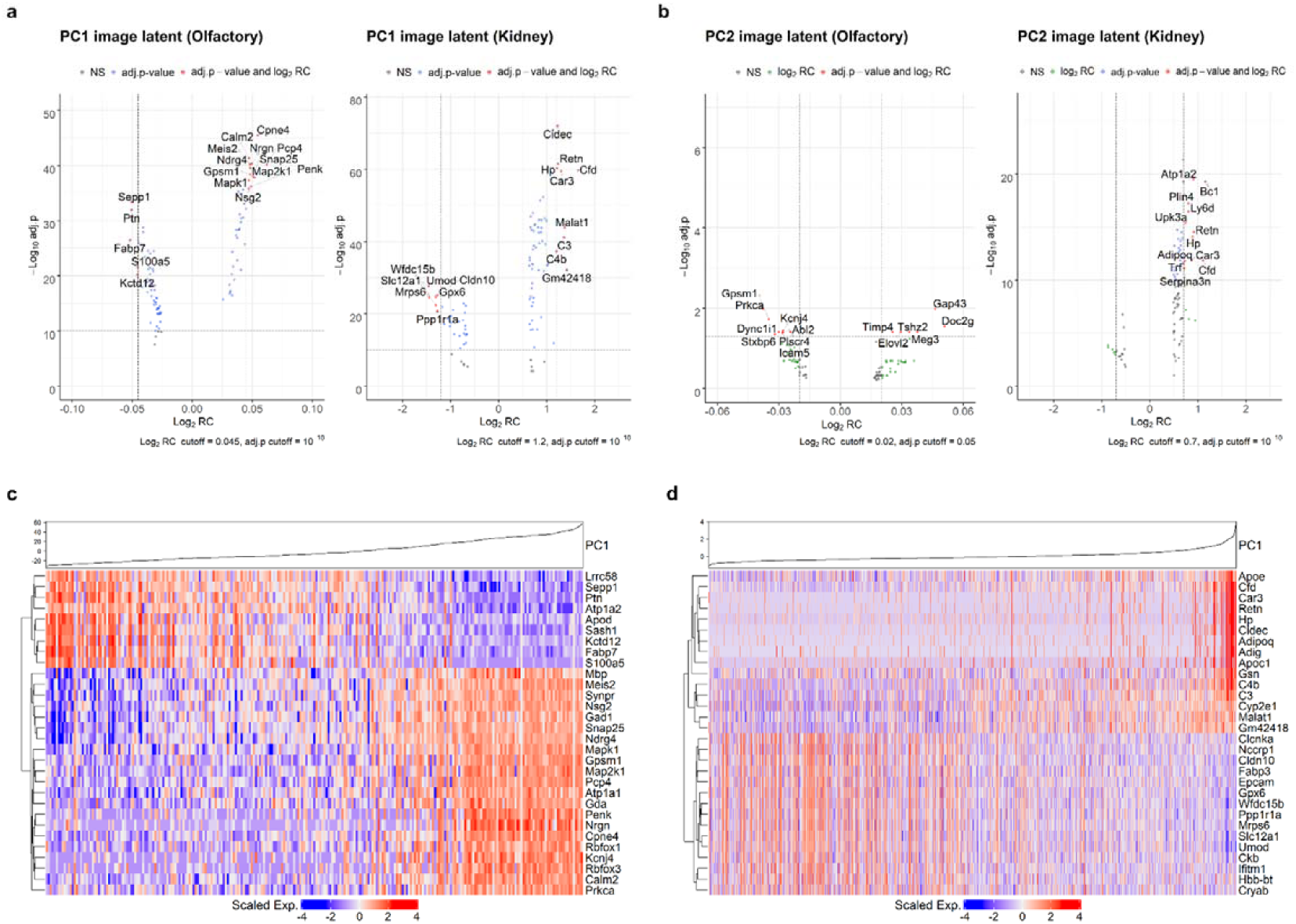
Investigation of morphological marker genes in olfactory bulb and kidney data. a, Volcano plots for highly associated genes with PC1 image latent features from olfactory and kidney tissues. Cutoff for log_2_ regression coefficient (RC) is 0.045 in olfactory bulb and 1.2 in kidney tissue. Cutoff for adjusted p-value (Benjamini-Hochberg correction) is 10^−10^ for both tissues. b, Volcano plots for highly associated genes with PC2 image latent features from olfactory bulb and kidney data. Cutoff for log_2_RC is 0.02 and 0.7 and cutoff for adjusted p-value (Benjamini-Hochberg correction) is 0.05 and 10^−10^, respectively. c, Heatmap for top 30 highly associated genes for log_2_RC in PC1 image latent space from olfactory bulb tissue. Hierarchical clustering was performed for top 30 genes and PC1 value in each of the spot was shown on top d, Heatmap for top 30 highly associated genes for log_2_RC in PC1 image latent space from kidney tissue. Hierarchical clustering was performed for top 30 genes and PC1 value in each of the spot was presented on top.

Top 5 genes for PC1 were *PCP4, CPNE4, MAP2K1, FABP7*, and *SNAP25* in olfactory bulb and *CFD, WFDC15B, UMOD, SLC12A1*, and *GM42418* in kidney tissue. Among the top 5 markers genes in olfactory bulb, *FABP7* and *DOC2G* exhibited variable expression across five distinct layers and are known markers for glomerular and mitral cell layer, respectively. (**Fig. 4a, b**)^18^. Also in the kidney tissue, the marker genes showed spatially varying patterns of expression in renal capsule, cortex and medulla (**Fig. 4c, d**). Unlike breast tissue, the genes associated with PC1 and PC2 were differently extracted in both the tissues.

**Fig. 4:**
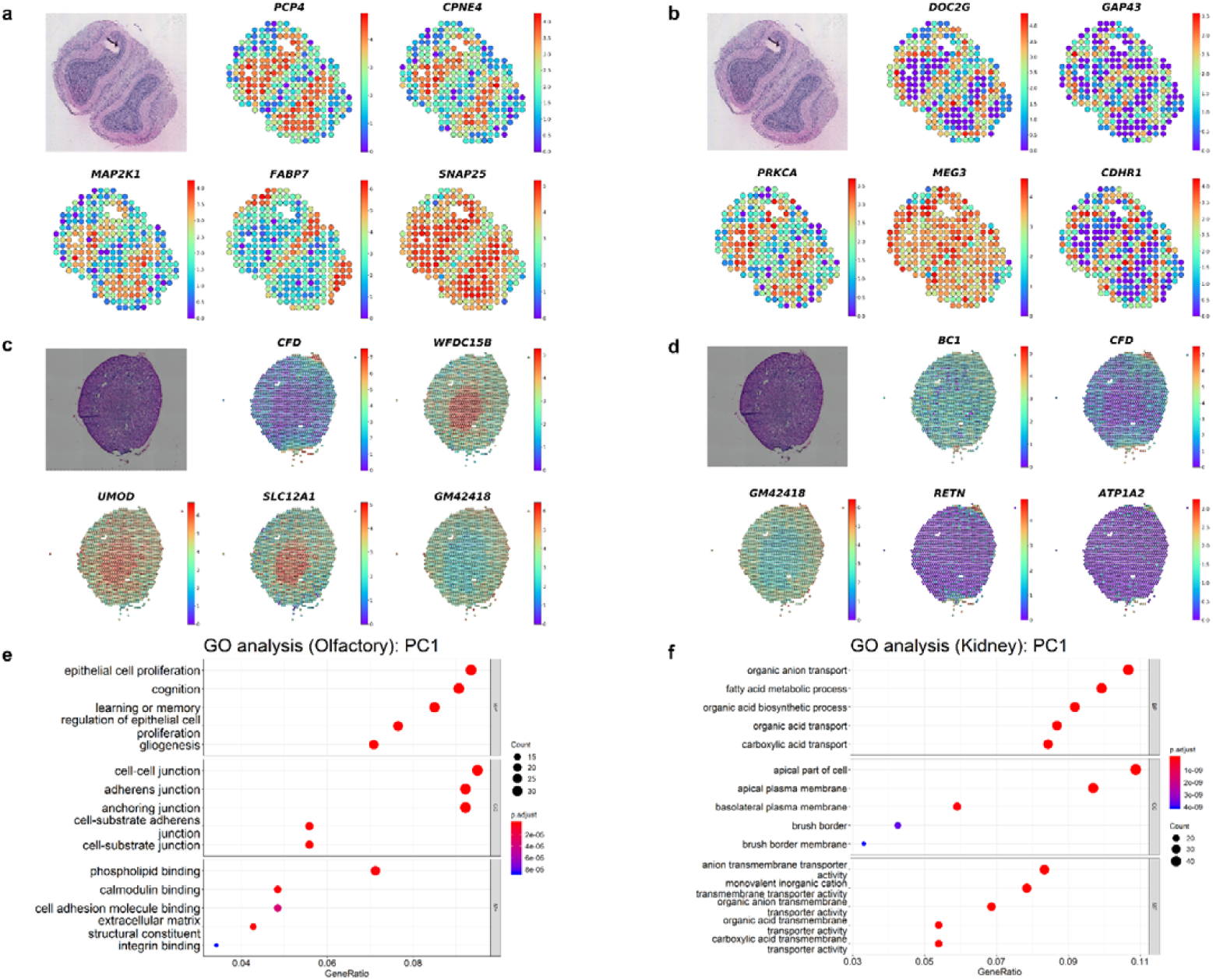
Spatial gene expression pattern and functional aspect of SPADE in olfactory bulb and kidney data. a, Spatial expression of top 5 genes representing greatest contrast in PC1 image latent space in olfactory bulb tissue. b, Spatial expression of top 5 genes representing greatest contrast in PC2 image latent space from olfactory bulb tissue. c, Spatial expression of top 5 genes representing greatest contrast in PC1 image latent space from kidney tissue. d, Spatial expression of top 5 genes representing greatest contrast in PC2 image latent space from kidney tissue. e, Gene ontology (GO) analysis for SPADE genes from PC1 image latent in olfactory bulb data. Top 10 GO terms for each subcategory, molecular function (MF), cellular component (CC), and biological process (BP) were exhibited. Number of overlapped genes was expressed as size of dot and Benjamini-Hochberg adjusted p-value was exhibited with colormap f, GO analysis for SPADE genes from PC2 image latent in kidney data. Top 10 GO terms for each subcategory, MF, CC, and BP were exhibited. Number of overlapped genes was expressed as size of dot and Benjamini-Hochberg adjusted p-value was exhibited with colormap

### Functional molecular terms and clustering based on SPADE for olfactory bulb and kidney tissues

Functional gene enrichment analysis showed over-represented biological process GO terms including epithelial cell proliferation and cognition in PC1 and PC4 SPADE genes of olfactory bulb (**Fig. 4e, Supplementary Fig. 10**). GO terms of fatty acid metabolic process or organic acid biosynthesis were identified from PC1 to PC4 in kidney tissue (**Fig. 4f, Supplementary Fig. 10**). For the molecular function, terms such as phospholipid and calmodulin binding is enriched in olfactory bulb while transmembrane transporter activity in kidney tissue.

Spot clustering was performed based on SPADE genes from PC1 and PC2 image latents and clusters were separated on t-SNE plots of transcriptomic data and image latents (**Fig. 5a, b**). Marker genes of each SPADE-based cluster were extracted (**Supplementary Fig. 11a, b**). Meanwhile, conventional spot clustering with HVG and marker discovery was performed and the clusters were presented on t-SNE plot (**Supplementary Fig. 12**). Heatmaps for SPADE genes showed distinguishable gene expression patterns across different SPADE-based and HVG-based clusters (**Supplementary Fig. 13**). The mismatched clusters were also identified in both olfactory bulb and kidney tissues. (**Fig. 5c, d**). The mapping of SPADE and HVG-based clusters on the tissue exhibited spatial distribution of the clusters according to the image features (**Fig. 5e, f**). In the olfactory bulb data, the greatest number of mismatched spots were observed in SPADE 4-HVG 1 cluster (69.57%; 16/23). All of the spots were located at the boundary of glomerular layer (mainly cluster 1) and olfactory nerve layer (mainly cluster 4) (**Fig. 5g**). When the SPADE4-HVG1 mismatched cluster was mapped on image latent PC plot, the spots were closer to SPADE 4-HVG 4 (median distance: 13.98) spots than SPADE 1-SPADE 1 spots (median distance: 19.35) (p=0.067) (**Fig. 5h**). In kidney tissue, SPADE 2-HVG 1 was the top mismatched spots, distributed on the renal capsule or cortical layers (**Supplementary Fig. 14a**). SPADE 2-HVG 1 spots were not significantly closer to SPADE 2-HVG 2 (median distance: 0.59) than SPADE 1-HVG 1 (distance: 0.61) (p=n.s.) (**Supplementary Fig. 14b**).

**Fig. 5:**
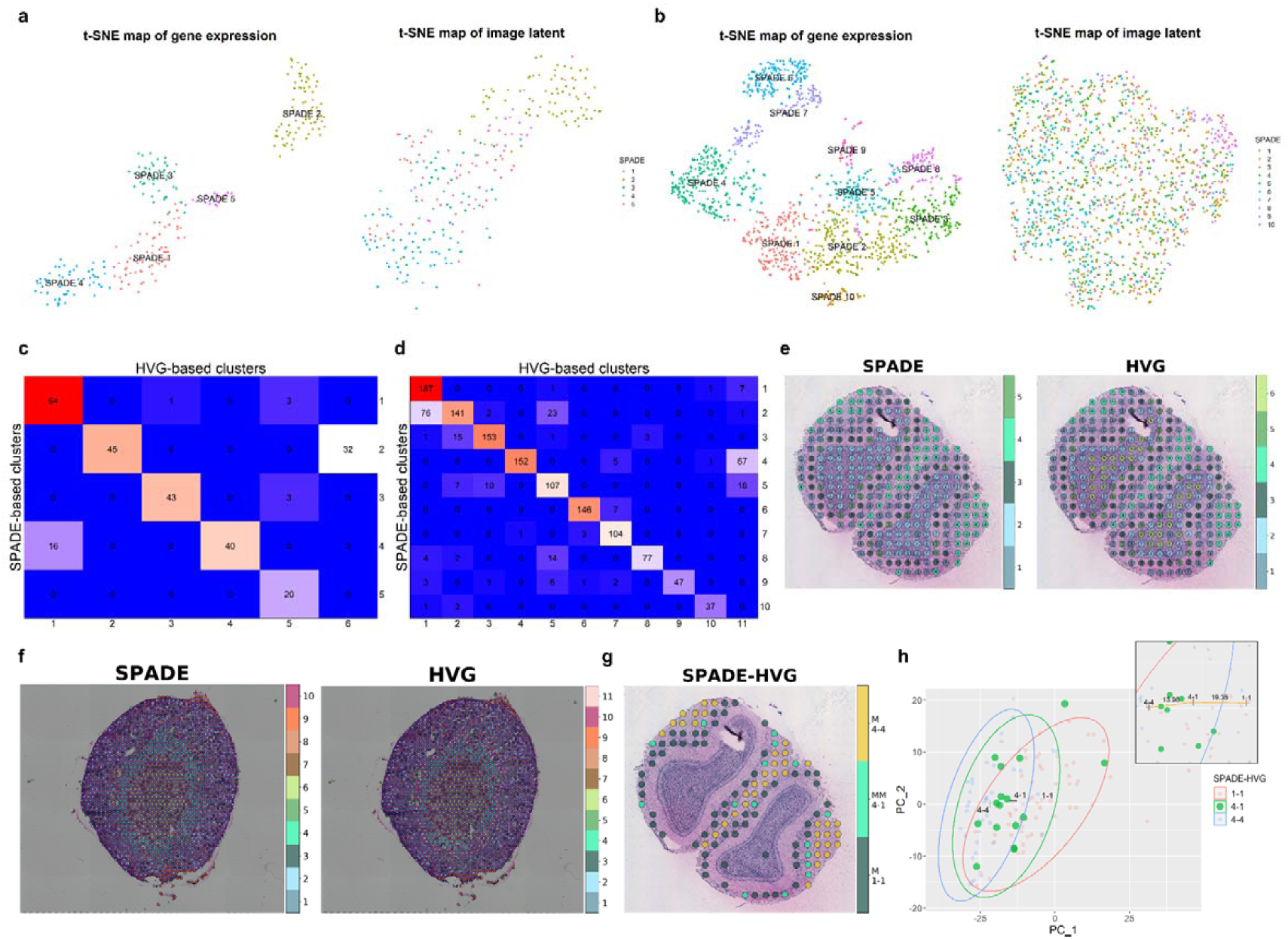
Spot clustering based on SPADE genes in olfactory bulb and kidney data. a, t-SNE plot of transcriptomic data and deep learning-derived image latent features in olfactory bulb tissue. SPADE genes were utilized to classify spots into 5 SPADE-based clusters and the cluster identity of each spot was visualized. b, t-SNE plot of transcriptomic data and deep learning-derived image latent features in kidney tissue. SPADE genes were utilized to classify spots into 10 SPADE-based clusters and the cluster identity of each spot was visualized. c, A cross table exhibiting number of spots overlapped in corresponding SPADE and HVG-based clusters in olfactory bulb tissue. d, A cross table exhibiting number of spots overlapped in corresponding SPADE and HVG-based clusters in kidney tissue. e, Spatial distribution of SPADE and HVG-based spot clusters mapped on the olfactory bulb tissue. The cluster numbers for SPADE or HVG-based cluster were exhibited on the right panel. f, Spatial distribution of SPADE and HVG-based spot clusters mapped on the kidney tissue. The cluster numbers for SPADE or HVG-based cluster were exhibited on the right panel. g, Spatial mapping of the SPADE 1-HVG 1 and SPADE 4-HVG 4 matched and SPADE 4-HVG 1 mismatched spot clusters in olfactory bulb tissue. M is abbreviation for matched cluster and MM is for mismatched cluster. h, PC1 and PC2 image latent plot for matched and SPADE 4-HVG 1 mismatched spot clusters in olfactory bulb tissue. A distribution of the mismatched spot clusters was presented in green dots. 95% confidence ellipse for each cluster based on multivariate t-distribution was exhibited on PC plot. Enlarged picture on the top shows median distance between mismatched spots and center of mass for matched clusters.

## Discussion

SPADE which integrates histological image patterns with spatial gene expression data discovered genes associated with morphological landscape. The analysis on three datasets across two different spatial transcriptomics (ST) platforms showed a flexible and scalable application of SPADE to various platforms as well as tissue types. The mapping of SPADE genes on tissue images showed biologically crucial spatial patterns that distinguish the morphological architectures. Moreover, SPADE was able to analyze biological processes related to morphological heterogeneity of tissues by using conventional analytic methods such as gene ontology. Since SPADE genes were related to heterogeneous image patterns, it was suggested that clustering based on these genes can well preserve morphological contexts.

SPADE can be applied as an explorative tool to unveil key genes or enriched biological process related to spatial and morphological heterogeneity of tissue. In breast cancer tissue, *MALAT1* gene showed the greatest variation of expression associated with PC1 image latent feature. *MALAT1* is highly expressed in cancer tissues and is related to cancer cell proliferation via ERK/MAPK pathway^19^. As *MALAT1* explained spatial and cellular heterogeneity of tumor tissue, it can be assumed as a key molecular feature underlying the heterogeneity of tumor microenvironment. Furthermore, the enriched GO terms for SPADE genes of breast cancer tissue included extracellular matrix and lymphocyte and leukocyte activation which are important components of the tumor microenvironment, exhibited under H&E staining^20^. The biological implications of SPADE genes were also found in olfactory bulb and kidney tissues. *PCP4*, one of the spatial marker genes in olfactory bulb (**Fig. 4a**) and a gene functioning as calmodulin binding protein, is known to be highly expressed in olfactory bulb and present different level of protein expression in granule cell and mitral cell layer^21^. Meanwhile, *SLC12A1*, also known as NKCC2 (Na-K-Cl cotransporter 2) is specifically expressed on thick ascending limb of Henle and located mainly in apical part of the cell^22^. *SLC12A1* was selected as a spatial marker in kidney tissue (**Fig. 4**) and among the GO terms, apical part of cell and anion transmembrane transporter activity were highly enriched in SPADE genes of kidney tissue. Besides, fatty acid oxidation which is one of the highly enriched GO terms, is an important process for ATP production especially in kidney proximal tubule^23^.

The molecular functions identified by SPADE showed variable functional enrichment patterns according to different image latent features. In case of breast cancer tissue, PC3 and PC6 explained 8.47% and 1.53% of variance of image features and were associated with GO terms such as response to metal ion and response to hypoxia, respectively. It has been reported that metal ions along with glycosaminoglycan in extracellular matrix affect angiogenesis and cancer cell migration via integrin activation^24^. In addition, hypoxia play a key role in tumor progression by hypoxia inducible factor (HIF) pathway and promotes angiogenesis and epithelial-to-mesenchymal transition^25^. In olfactory bulb tissue, image latents PC3 and PC4 were associated with GO terms such as calmodulin binding. Neurogranin, a protein that binds to calmodulin and modifies downstream signaling pathway in neurons, is localized in granule cell layer of olfactory bulb^26^. As different molecular functions were discovered according to the image latent features, SPADE could lead to the analysis of close interaction of molecular function and morphological patterns.

One of the applications of SPADE is clustering of spots by preserving morphological landscape. Since SPADE genes are representative genes responsible for morphological features, clustering of spots based on these genes could reflect the variability of image-level patterns. Accordingly, the mismatched spots were morphologically closer to SPADE than HVG-based clusters as presented by the distance between the clusters in the image latent space (**Fig. 2g, Supplementary Fig. 6, and Fig 5g, h**). This clustering method could be used to identify tissue architectures such as brain cortical layers with unsupervised manner, because this type of architectures are originally defined by morphological features^27^.

There are several factors that may have influenced the SPADE analysis. Since image patches corresponding to spots are provided as input, the density of the spots and size of image patches can significantly affect the result. Distance between center of spots in human breast and mouse kidney data from 10x genomics was approximately 100 micrometers while olfactory bulb dataset was 200 micrometers^4^. Detailed morphological information could have been lost due to sparse distribution of spots in olfactory bulb tissue and value of SPADE in exploring marker genes may have been underestimated. As recent progress in acquisition methods of spatial transcriptomic data can provide gene expression information with higher spatial resolution^5^, optimized parameters such as the size of image patches will be needed.

Our approach can provide an insight into the close relationship between molecular function and structure by identifying important genes responsible for the morphological landscape. A few methods have been proposed to find spatially variable genes employing location information of spots and focusing on the representative patterns of spatial gene expression^28-30^. However, they did not employ image features, thus, these methods identify spatially variable markers instead of markers associated with morphologic features. Another analysis tool, SpaCell, utilized spatial transcriptomic data along with tissue image features derived from a deep neural network to identify cell type and classify stage of disease^31^. While SpaCell applied the deep learning model to classify cell type and disease stage labels with a supervised manner, feature extraction with SPADE provides unbiased information about morphologically important genes associated with the histological features. Furthermore, SPADE can be combined with different types of images such as noninvasive imaging and immunohistochemistry which provides expression patterns of specific protein if spatially co-registered images are available. In other words, by using different types of images, transcriptomes associated with spatial patterns of a specific protein or function can be analyzed to obtain key markers and investigate functional interactions.

In conclusion, SPADE is flexibly used to interrogate molecular profiles responsible for tissue morphological landscape by listing up important genes and biological processes. The integration of different types of data, image and spatially resolved transcriptome, can provide an insight into the close relationship between structure and molecular functions, which eventually leads to comprehensive explanation of the pathophysiology of various diseases.

## Methods

### Data

H&E stained slides and count data of gene expression for the spots of human breast cancer tissue and mouse kidney tissue were obtained from publicly available dataset provided by 10x Genomics (https://www.10xgenomics.com/resources/datasets/). For human breast cancer, ‘Block A section 1’ data which contains total 3,813 spots and 33,538 genes was used for the analysis. Tissue slide images, scale factors, coordinates of spots and count data were used for the analysis. Mouse kidney tissue data contains total 1,438 spots and 31,053 genes. High resolution and low resolution images, scale factors, coordinates of spots and UMI count data were included in the study.

A slide image of mouse olfactory bulb tissue and count data of gene expression for the spots were obtained from publicly available dataset provided by SciLife laboratory (https://www.spatialresearch.org/resources-published-datasets/). Among the 12 section slides of olfactory bulb, ‘MOB Replicate 1’ which contains 267 spots and 16,383 genes was utilized for the further analysis. Tissue slide images, spot coordinates and transformation matrix were downloaded.

### Image feature extraction from tissue images

A high resolution H&E stained slide was cropped into multiple square patches. The patch size depends on the size of the entire tissue image. The size of the image in breast cancer, olfactory bulb, and kidney tissues was 2,000×2,000, 9,931×9,272, and 2,000×2,000 pixels and patch size was 32×32, 400×400, and 32×32 pixels, respectively. The center of the image patch was determined by sampling spot coordinates from the ST data. Each image patch was provided as an input for a pre-trained CNN (**Fig. 1a**). As a pretrained CNN model, we used VGG-16, which was trained by the classification task of natural images of ImageNet data^12, 32^. The VGG-16 model was used as a feature extractor. Thus, the last layer which consists of 1,000 nodes for classification labels for ImageNet challenge was removed. In addition, to apply patch-based approach which can have variable patch size according to the size of entire tissue image, convolutional-only layers were included. The last convolutional layer produced two-dimensional images instead of vectors, thus, global-average pooling layer was added considering size-adaptive feature extractor^33^. This feature extractor produced 512-dimensional vectors.

### Dimension reduction for image features

To visualize image features of all patches corresponding to spots, t-distributed stochastic neighbor embedding (t-SNE) was employed^17^. t-SNE is a non-linear method of reducing dimensions of data and visualizing high-dimensional data in low-dimensional space. The perplexity was set at 30 and initialization of embedding was based on principal component analysis (PCA).

PCA was performed to reduce the dimension of 512 features obtained from the VGG-16 model and extract principal components (PCs). These PCs were also mapped on the tissue image according to the location of patches to visualize spatial distribution patterns of image features. The whole process was implemented in Python version 3.7.0 and scikit-learn (ver. 0.21.3.).

### SPADE genes

A function *SCTransform* in R package Seurat (version 3.1.4) was applied to normalize feature counts in each spot and to find top 1,000 highly variable genes (HVG), which show variability of expression across spots^34^. A linear model was generated to fit scaled gene expression of HVG to PCs of image latent features. Empirical Bayes algorithm in R package limma (version 3.42.2) was applied and associated genes based on linear regression analysis for the value of PCs were ranked according to regression coefficient (RC) or corrected p-value with Benjamini-Hochberg method (**Fig. 1a**)^35^. Results for linear regression analysis for PC1 and PC2 were visualized by R package EnhancedVolcano (version 1.4.0)^36^.

List of the genes presenting false discovery rate (FDR) less than 0.05 in PC1, PC2, PC3, PC4, PC5, or PC6 were gathered to select SPADE genes. SPADE genes were filtered by log RC over 0.25 for breast and kidney tissue data and over 0.01 for olfactory bulb data. The threshold was determined according to the results of limma considering number of SPADE genes. For spot clustering, genes derived from PC1 and PC2 were pooled and filtered to adjust number of the SPADE genes between 300 and 500. The whole process was performed in R version 3.6.1.

### Gene ontology analysis

Gene ontology (GO) analysis was implemented with R package clusterProfiler (version 3.14.3) using *enrichGO* function^13, 14, 37^. Top 10 enriched GO terms in subcategories including biological process (BP), cellular component (CC) and molecular function (MF) were extracted based on SPADE genes from PC1 to PC6 image features (**Fig. 2a and Fig. 4e, f**). P-value was corrected with Benjamini-Hochberg false discovery rate and the cutoff was 0.05.

### Clusters based on expression data of selected genes

The sampling spots were conventionally clustered according to scRNA-seq analysis workflow in R package Seurat^38^. Spatial information of each spot was not included in this clustering process. *SCTransform* was performed and 1000 HVG were selected. Dimension reduction with PCA followed by shared nearest neighbor (SNN) graph-based spot clustering using Louvain algorithm were done^15, 16^. DEG analysis for each highly variable gene (HVG)-based cluster was performed by *FindAllMarkers* and marker genes were visualized by a heatmap. As another clustering based on SPADE, SPADE genes were used to obtain a SNN graph of spots. Dimension reduction with PCA on SPADE genes followed by SNN graph-based spot clustering using Louvain algorithm were performed as HVG. SPADE-based cluster number was rearranged so that *n*-th SPADE-based cluster shared the most of spots with *n*-th HVG-based cluster. Marker genes for each SPADE-based cluster were also extracted by *FindAllMarkers* and visualized by a heatmap.

The expression of SPADE genes and clusters were visualized using ComplexHeatmap (version 2.2.0)^39^. The clusters generated from SPADE genes and from HVG were compared. In addition, SPADE and HVG-based clusters were spatially mapped on tissue image (**Fig. 2f and Fig. 5e, f**). Mismatched spot clusters which have different SPADE and HVG-based cluster number (*n* and *m*, respectively) were mapped on the tissue image (**Fig. 2g and Fig. 5g**) and also on the image latent space represented by PCs of image features (**Fig. 5h**). The center of mass (COM) for matched clusters, *n*-th SPADE-*n*-th HVG and *m*-th SPADE-*m*-th HVG, in 512-dimensional image latents was calculated. Euclidean distances between spots comprising the mismatched cluster and the COM of two matched clusters were computed. For the next step, the distance values were compared with Mann-Whitney U test. *P-values* below 0.05 were considered statistically significant.

## Data availability

Three publicly available dataset were utilized in this study. Human breast cancer and mouse kidney tissue data were downloaded from 10x genomics homepage (https://www.10xgenomics.com/resources/datasets/) while mouse olfactory bulb tissue data was from SciLife laboratory homepage (https://www.spatialresearch.org/resources-published-datasets/).

## Code availability

Python and R source code for SPADE is uploaded on https://github.com/mexchy1000/spade

## Acknowledgements

This research was supported by the National Research Foundation of Korea (NRF-2019R1F1A1061412, NRF-2019K1A3A1A14065446, and NRF-2020M3A9B6038086) and also supported by a grant of the Korea Health Technology R&D Project through the Korea Health Industry Development Institute (KHIDI), funded by the Ministry of Health & Welfare, Republic of Korea (HI19C0339).

## Author contributions

H.C. and D.S.L. designed the study. H.C. developed the model. S.B. collected and analyzed data. S.B. and H.C. modified the model. All authors contributed to the interpretation of the data and wrote the paper.

