## Supplementary Data for "Discovery of molecular features underlying morphological landscape by integrating spatial transcriptomic data with deep features of tissue image"

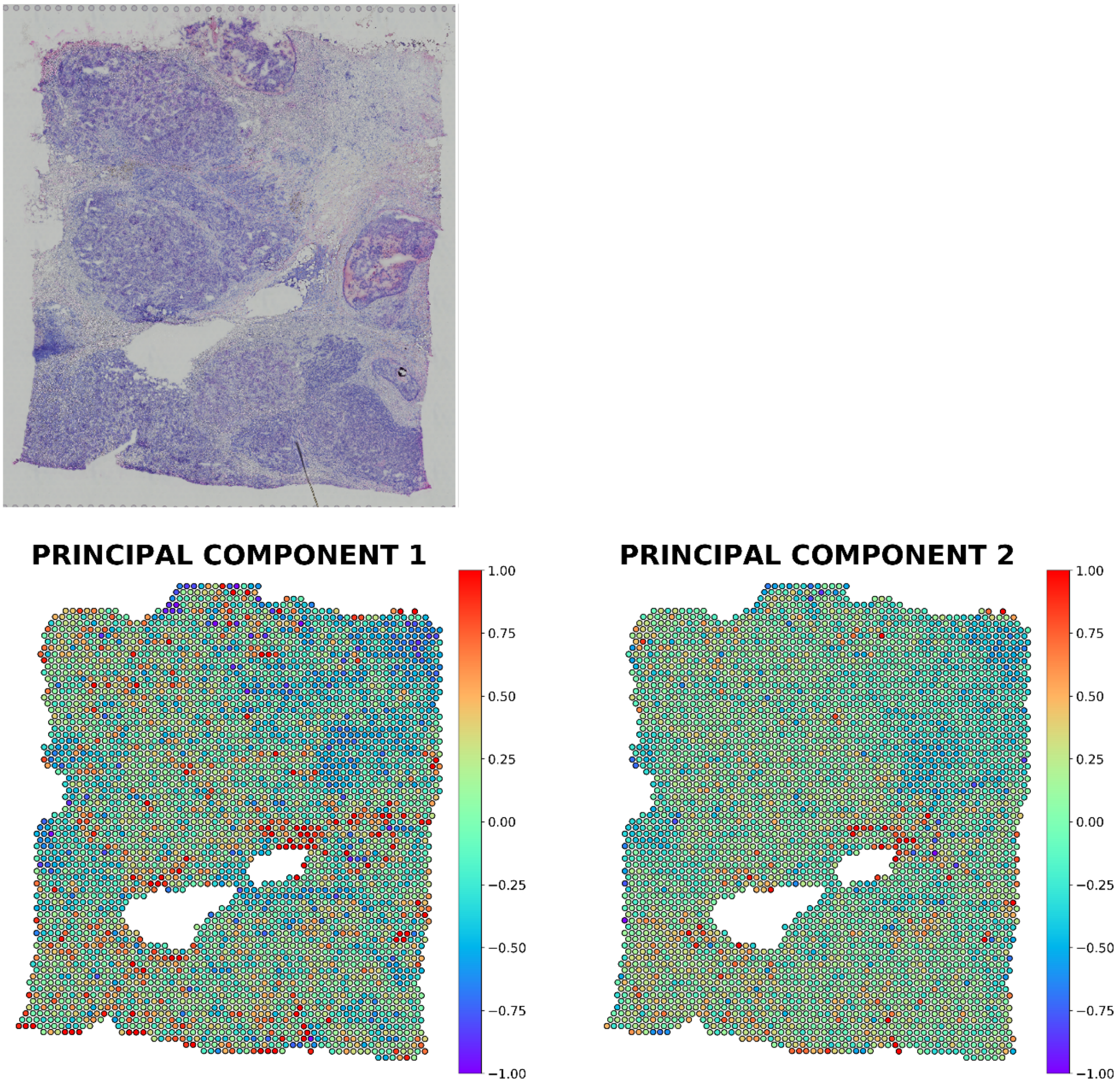


**Supplementary Figure 1. Spatial mapping of PC1 and PC2 image latent from breast cancer tissue.** PC1 or PC2 value of each spot was visualized using colormap.

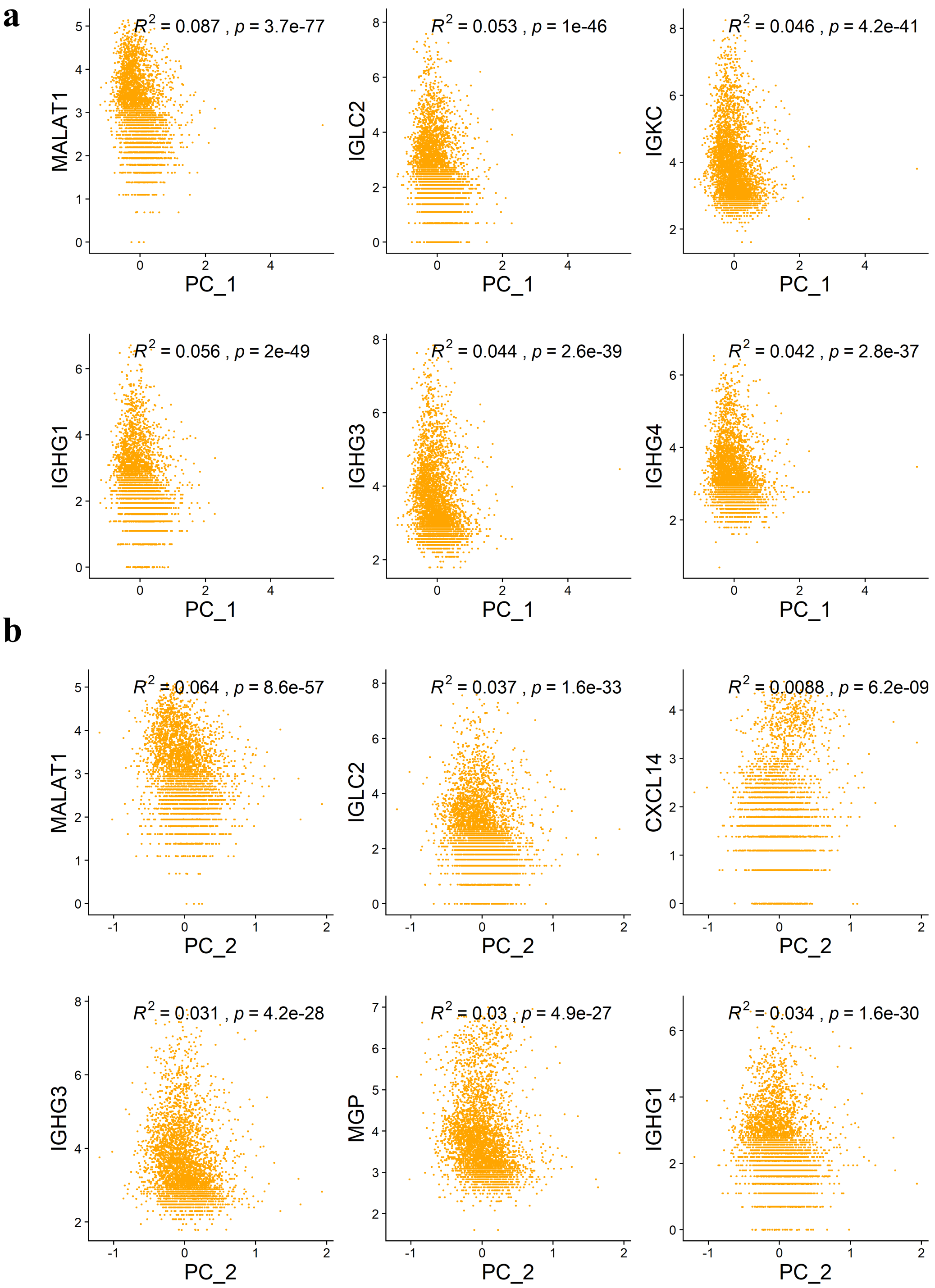


**Supplementary Figure 2. Top 6 genes associated with PCs of image latent features**

Scatter plot between (a) PC 1 or (b) PC 2 values and expression of top 6 highly associated genes for each PC image latent in breast cancer tissue. R-squared values and p-values for each scatter plot was exhibited on top of each plot.

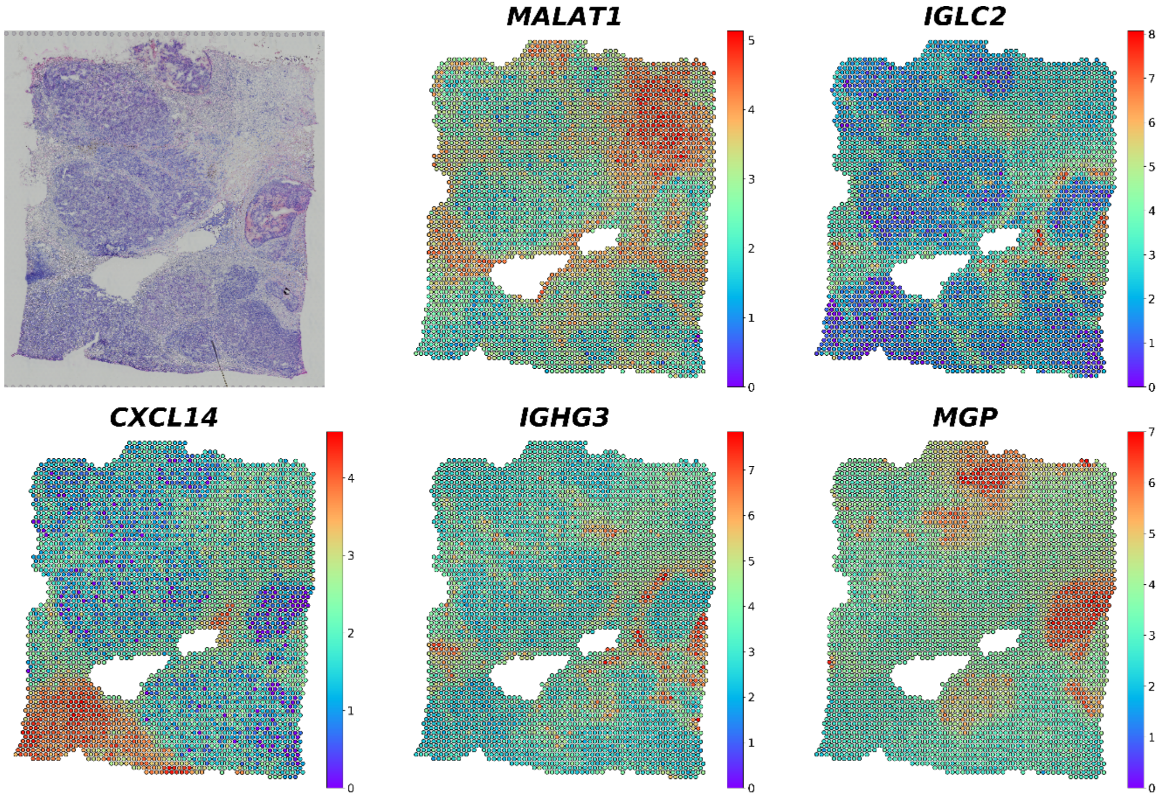


**Supplementary Figure 3. Spatial expression of top 5 genes representing greatest contrast in PC2 image latent space from breast cancer tissue.** Gene expression level in each spot was visualized with colormap.

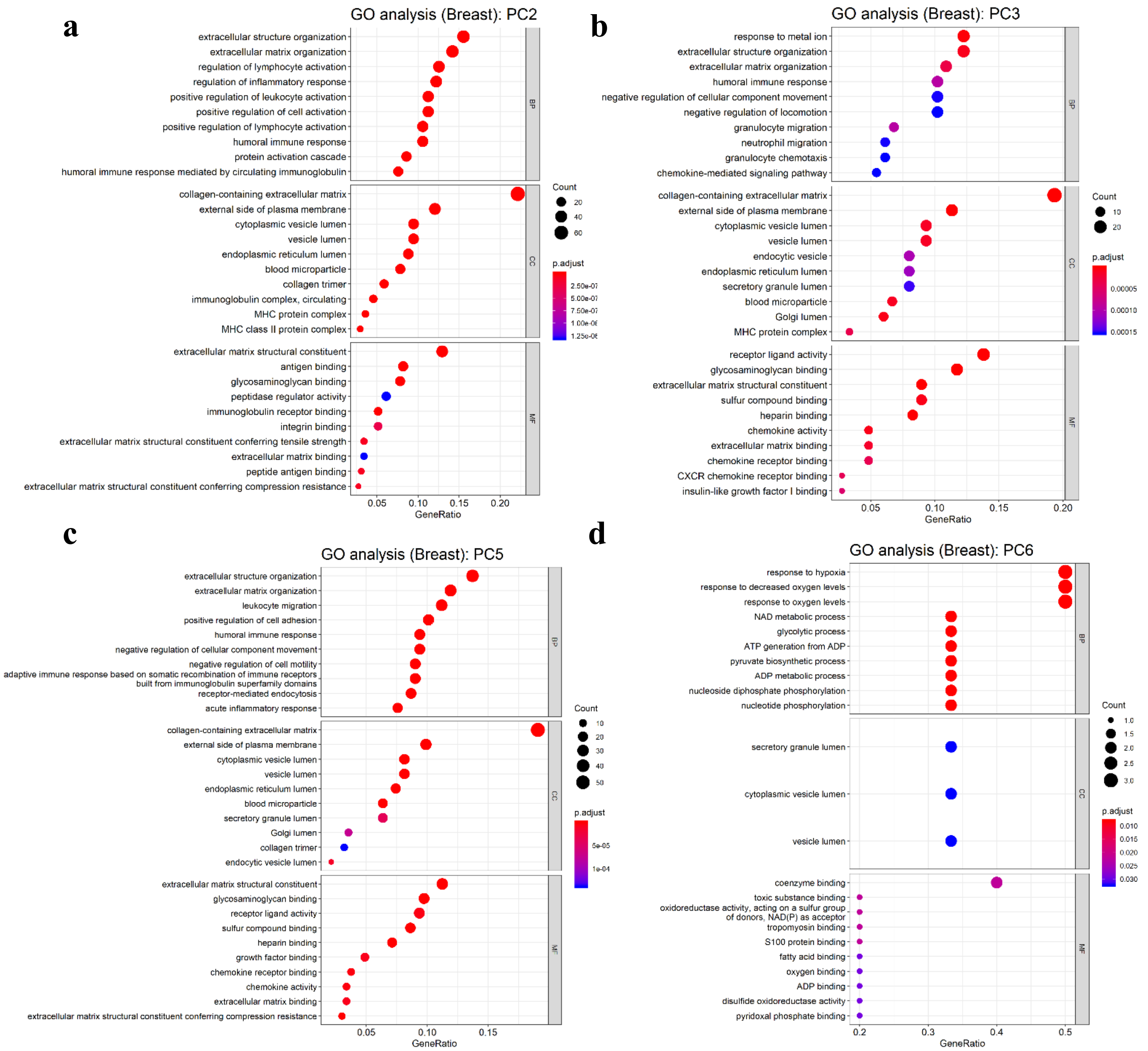


**Supplementary Figure 4. Gene ontology for SPADE genes extracted from different PCs.**

Gene ontology (GO) analysis for SPADE genes from (a) PC2, (b) PC3, (c) PC5, and (d) PC6 image latent in breast cancer tissue showing top 10 over-represented gene functions in each subcategory (MF: molecular function, CC: cellular component, BP: biological process). Number of overlapped genes was expressed as size of dot and Benjamini-Hochberg adjusted p-value was exhibited with colormap.

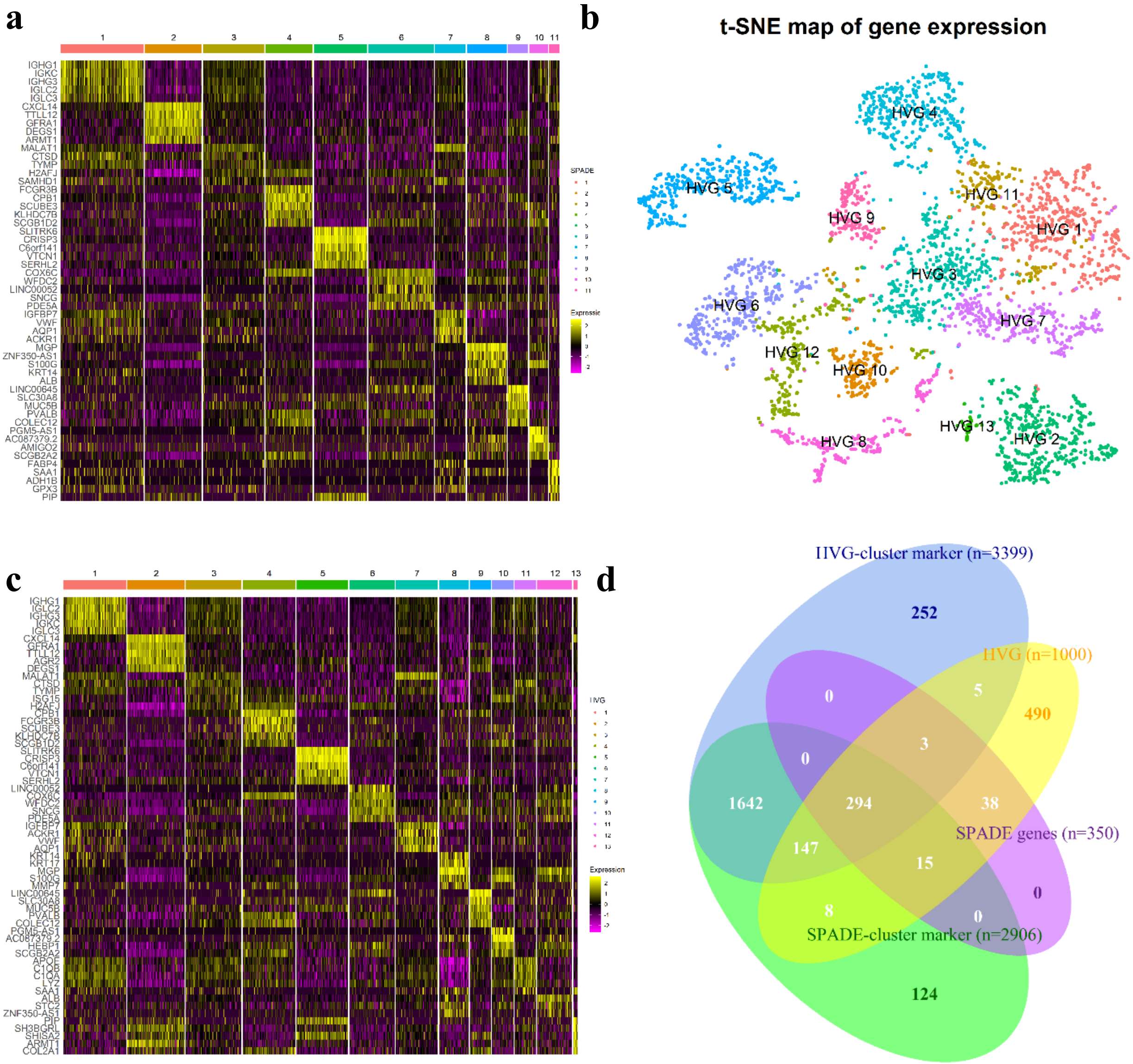


**Supplementary Figure 5. Spot clustering based on SPADE genes or HVG in breast cancer tissue** (a) Heatmap showing top 5 differentially expressed marker genes in each SPADE-based cluster. (b) t-SNE plot of transcriptomic data for HVG-based clusters. HVG were utilized to classify spots into 13 HVG-based clusters. (c) Heatmap showing top 5 differentially expressed marker genes in each HVG-based cluster. (d) Venn diagram showing the composition of SPADE-based cluster markers, HVG-based cluster markers, SPADE genes, and HVG. The number in the diagram represents number of spots corresponding to each category.

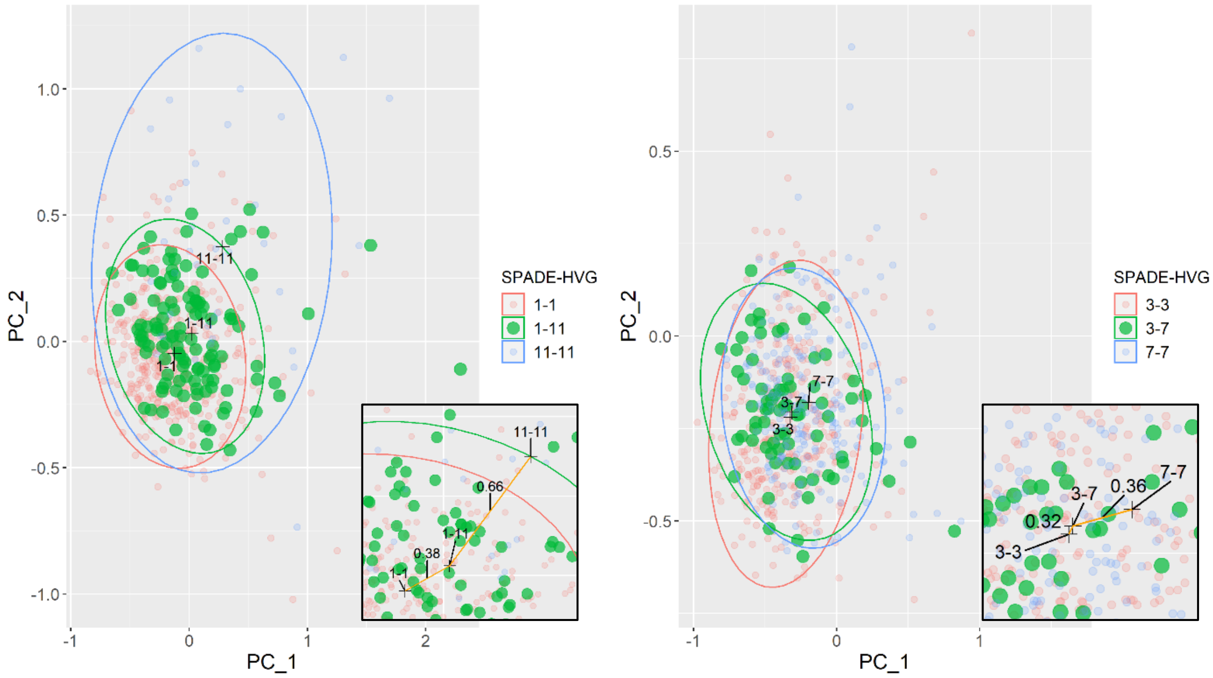


**Supplementary Figure 6. Image latents of mismatched clusters of different clustering methods.**

PC1 and PC2 image latent plot for matched and mismatched spot clusters. A distribution of mismatched spot clusters was presented in green dots. 95% confidence ellipse for each cluster based on multivariate t-distribution was exhibited on PC plot. Plot in the left shows SPADE 1-HVG 11 mismatched cluster and plot in the right shows SPADE 3-HVG 7 mismatched cluster. Enlarged pictures in the bottom show median distance between mismatched spots and center of mass for matched clusters.

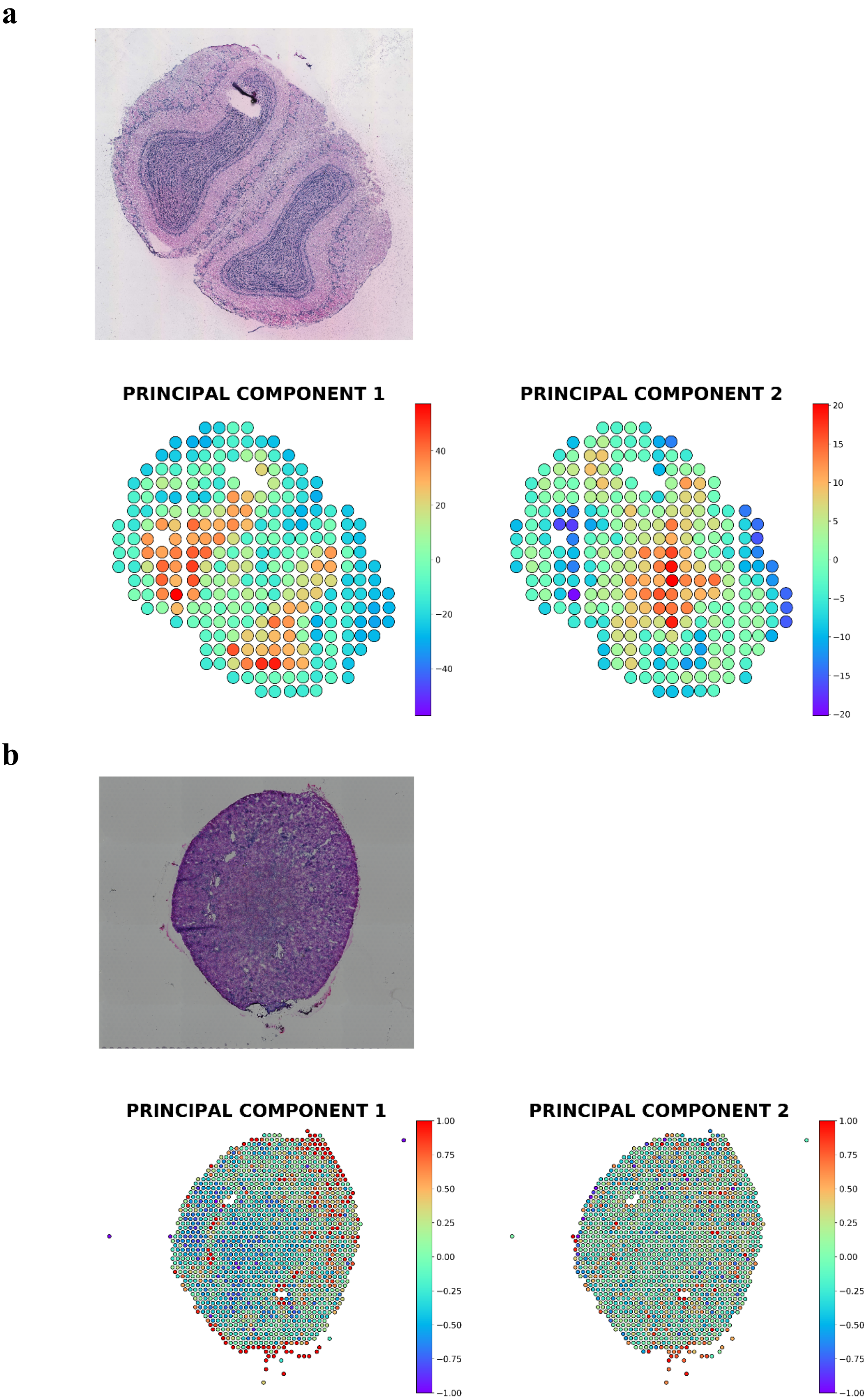
a
PRINCIPAL COMPONENT 1
40
20
0
−20
−40
PRINCIPAL COMPONENT 2
20
15
10
5
0
−5
−10
−15
−20
b
PRINCIPAL COMPONENT 1
1.00
0.75
0.50
0.25
0.00
−0.25
−0.50
−0.75
−1.00
PRINCIPAL COMPONENT 2
1.00
0.75
0.50
0.25
0.00
−0.25
−0.50
−0.75
−1.00

**Supplementary Figure 7. Spatial mapping of image latents in two different datasets.**

Spatial mapping of PC1 and PC2 image latent from (a) olfactory bulb and (b) kidney tissues. PC1 or PC2 value of each spot was visualized using colormap.

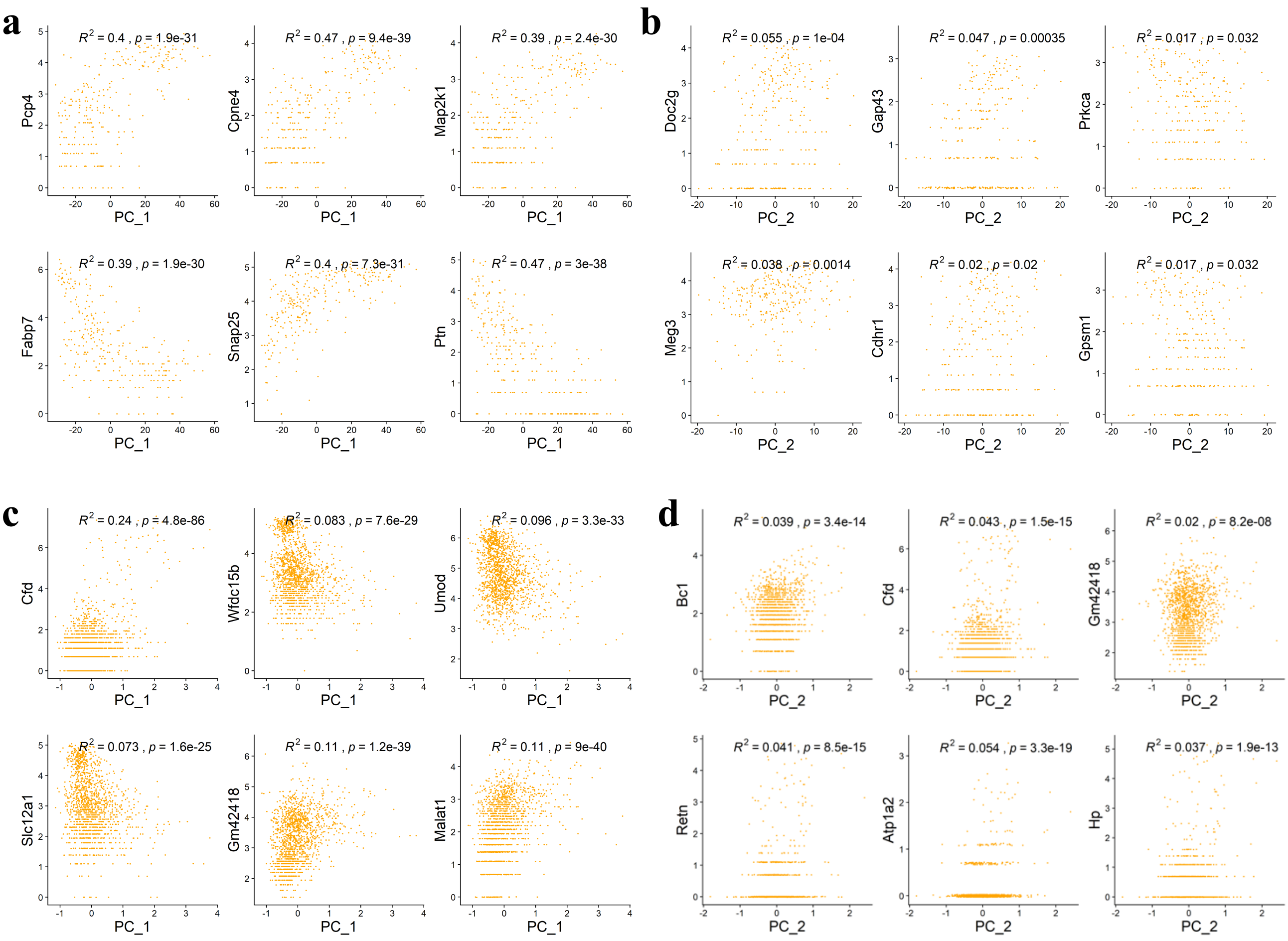


**Supplementary Figure 8. Top 6 associated genes with PC image latents.**

Scatter plot between PC values and expression of top 6 highly associated genes for each PC image latent. R-squared values and p-values for each scatter plot was exhibited on top of each plot. Plot for (a) PC1 or (b) PC2 value and top gene expression in olfactory bulb tissue. Plot for (c) PC1 or (d) PC2 value and top gene expression in kidney tissue.

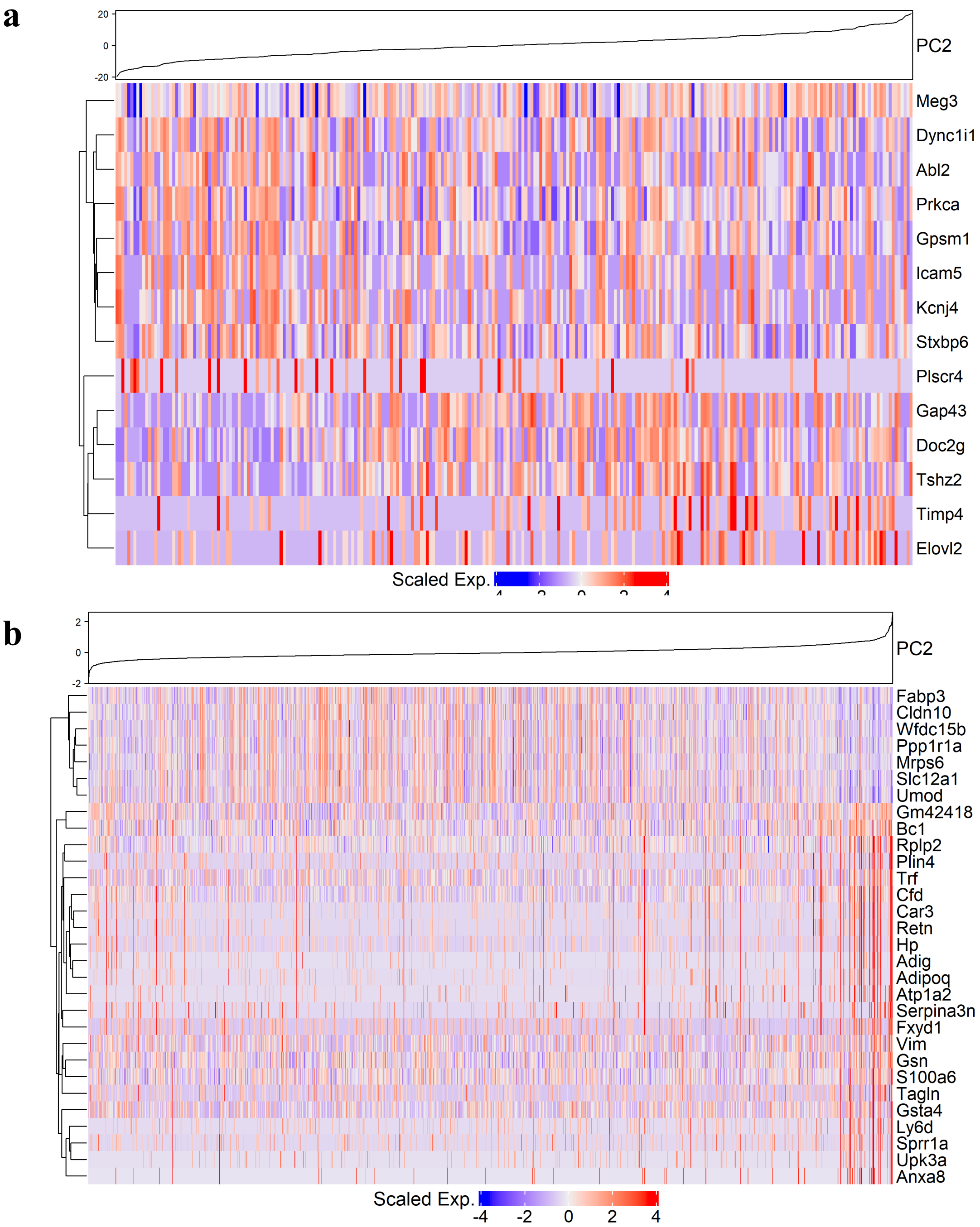


**Supplementary Figure 9. Heatmaps for genes associated with image latent PCs.**

Heatmap for (a) top 14 highly associated genes for $\log_2$ regression coefficient in PC2 image latent space from olfactory bulb and (b) top 30 highly associated genes from kidney data. Hierarchical clustering was performed for top 14 or 30 genes in row and PC2 value in each of the spot was shown on top.

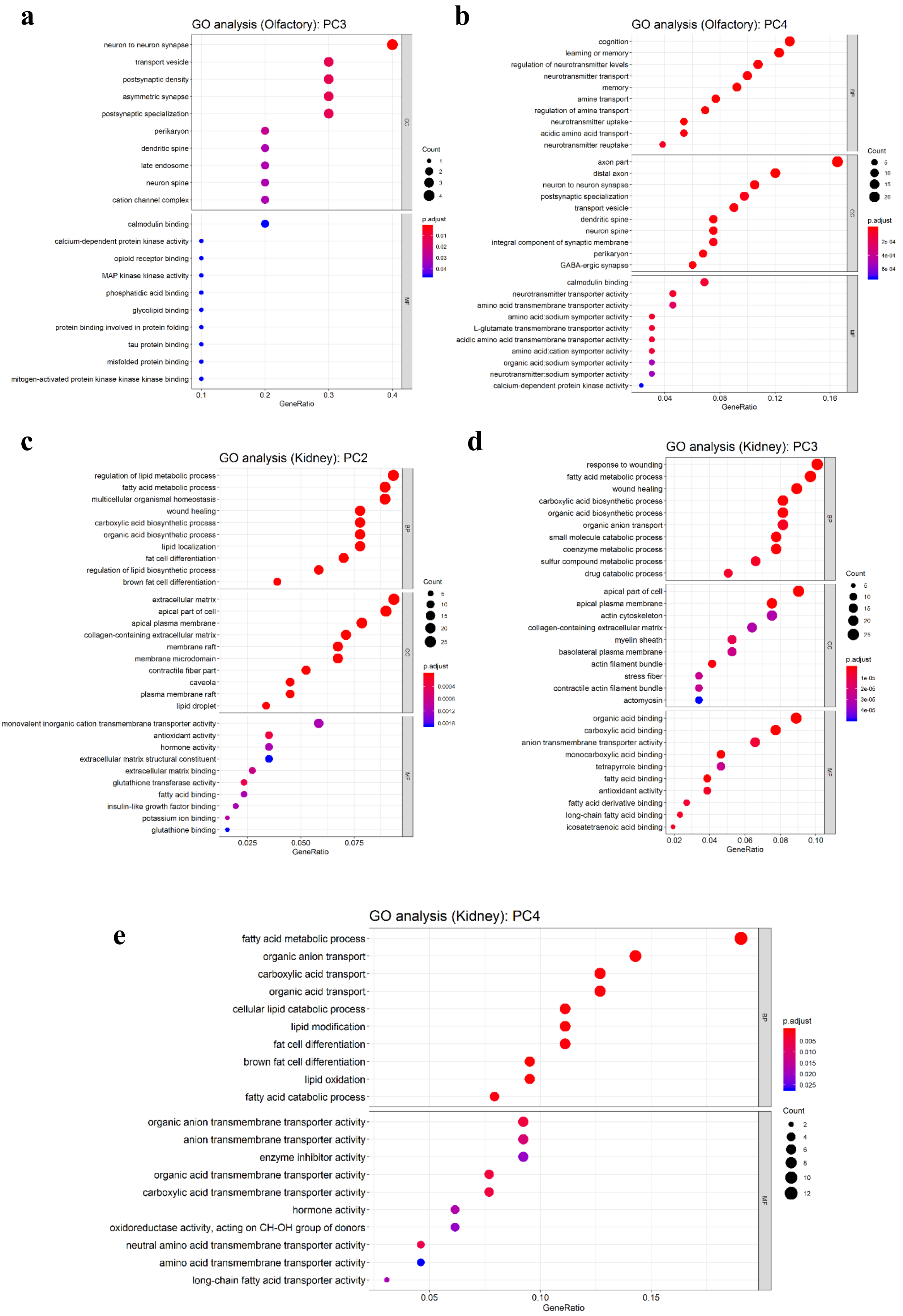
a
GO analysis (Olfactory): PC3
neuron to neuron synapse
transport vesicle
postsynaptic density
asymmetric synapse
postsynaptic specialization
perikaryon
dendritic spine
late endosome
neuron spine
cation channel complex
calmodulin binding
calcium-dependent protein kinase activity
opioid receptor binding
MAP kinase kinase activity
phosphatidic acid binding
glycolipid binding
protein binding involved in protein folding
tau protein binding
misfolded protein binding
mitogen-activated protein kinase kinase kinase binding
CC
MF
Count
p.adjust
GeneRatio
b
GO analysis (Olfactory): PC4
cognition
learning or memory
regulation of neurotransmitter levels
neurotransmitter transport
memory
amine transport
regulation of amine transport
neurotransmitter uptake
acidic amino acid transport
neurotransmitter reuptake
axon part
distal axon
neuron to neuron synapse
postsynaptic specialization
transport vesicle
dendritic spine
neuron spine
integral component of synaptic membrane
perikaryon
GABA-ergic synapse
calmodulin binding
neurotransmitter transporter activity
amino acid transmembrane transporter activity
amino acid:sodium symporter activity
L-glutamate transmembrane transporter activity
acidic amino acid transmembrane transporter activity
amino acid:cation symporter activity
organic acid:sodium symporter activity
neurotransmitter:sodium symporter activity
calcium-dependent protein kinase activity
BP
CC
MF
Count
p.adjust
GeneRatio
c
GO analysis (Kidney): PC2
regulation of lipid metabolic process
fatty acid metabolic process
multicellular organismal homeostasis
wound healing
carboxylic acid biosynthetic process
organic acid biosynthetic process
lipid localization
fat cell differentiation
regulation of lipid biosynthetic process
brown fat cell differentiation
extracellular matrix
apical part of cell
apical plasma membrane
collagen-containing extracellular matrix
membrane raft
membrane microdomain
contractile fiber part
caveola
plasma membrane raft
lipid droplet
monovalent inorganic cation transmembrane transporter activity
antioxidant activity
hormone activity
extracellular matrix structural constituent
extracellular matrix binding
glutathione transferase activity
fatty acid binding
insulin-like growth factor binding
potassium ion binding
glutathione binding
BP
CC
MF
Count
p.adjust
GeneRatio
d
GO analysis (Kidney): PC3
response to wounding
fatty acid metabolic process
wound healing
carboxylic acid biosynthetic process
organic acid biosynthetic process
organic anion transport
small molecule catabolic process
coenzyme metabolic process
sulfur compound metabolic process
drug catabolic process
apical part of cell
apical plasma membrane
actin cytoskeleton
collagen-containing extracellular matrix
myelin sheath
basolateral plasma membrane
actin filament bundle
stress fiber
contractile actin filament bundle
actomyosin
organic acid binding
carboxylic acid binding
anion transmembrane transporter activity
monocarboxylic acid binding
tetrapyrrole binding
fatty acid binding
antioxidant activity
fatty acid derivative binding
long-chain fatty acid binding
icosatetraenoic acid binding
BP
CC
MF
Count
p.adjust
GeneRatio
e
GO analysis (Kidney): PC4
fatty acid metabolic process
organic anion transport
carboxylic acid transport
organic acid transport
cellular lipid catabolic process
lipid modification
fat cell differentiation
brown fat cell differentiation
lipid oxidation
fatty acid catabolic process
organic anion transmembrane transporter activity
anion transmembrane transporter activity
enzyme inhibitor activity
organic acid transmembrane transporter activity
carboxylic acid transmembrane transporter activity
hormone activity
oxidoreductase activity, acting on CH-OH group of donors
neutral amino acid transmembrane transporter activity
amino acid transmembrane transporter activity
long-chain fatty acid transporter activity
BP
MF
p.adjust
0.005
0.010
0.015
0.020
0.025
Count
2
4
6
8
10
12
0.05
0.10
0.15
GeneRatio

**Supplementary Figure 10. Gene ontology analysis for SPADE genes of different datasets.**

Gene ontology (GO) analysis for SPADE genes from (a) PC3 and (b) PC4 image latent in olfactory bulb and (c) PC2, (d) PC3, and (e) PC4 image latent in kidney tissue. Top 10 GO terms for each subcategory, molecular function (MF), cellular component (CC), and biological process (BP) were exhibited. Number of overlapped genes was expressed as size of dot and Benjamini-Hochberg adjusted p-value was exhibited with colormap.

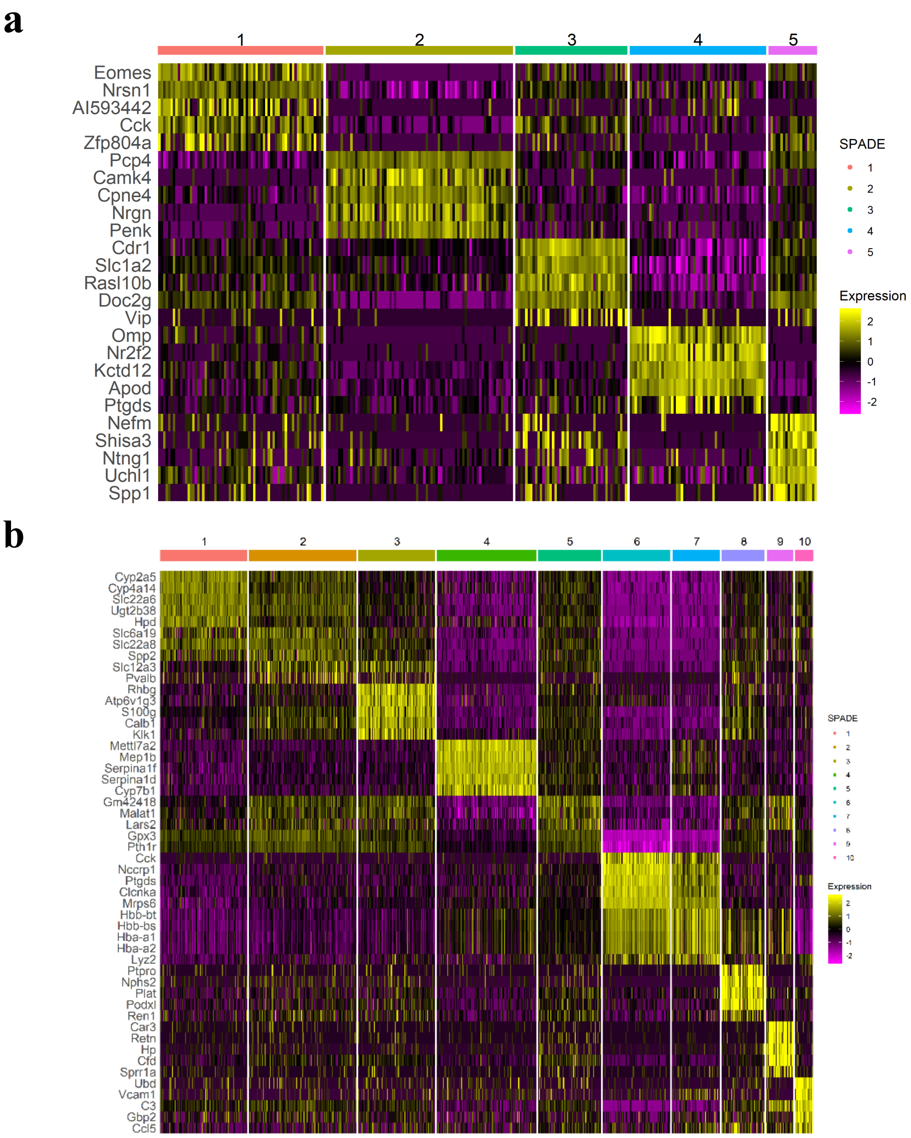


**Supplementary Figure 11. Heatmaps of markers for clustering**

Heatmaps showing top 5 differentially expressed genes in each SPADE-based cluster from (a) olfactory bulb and (b) kidney tissue.

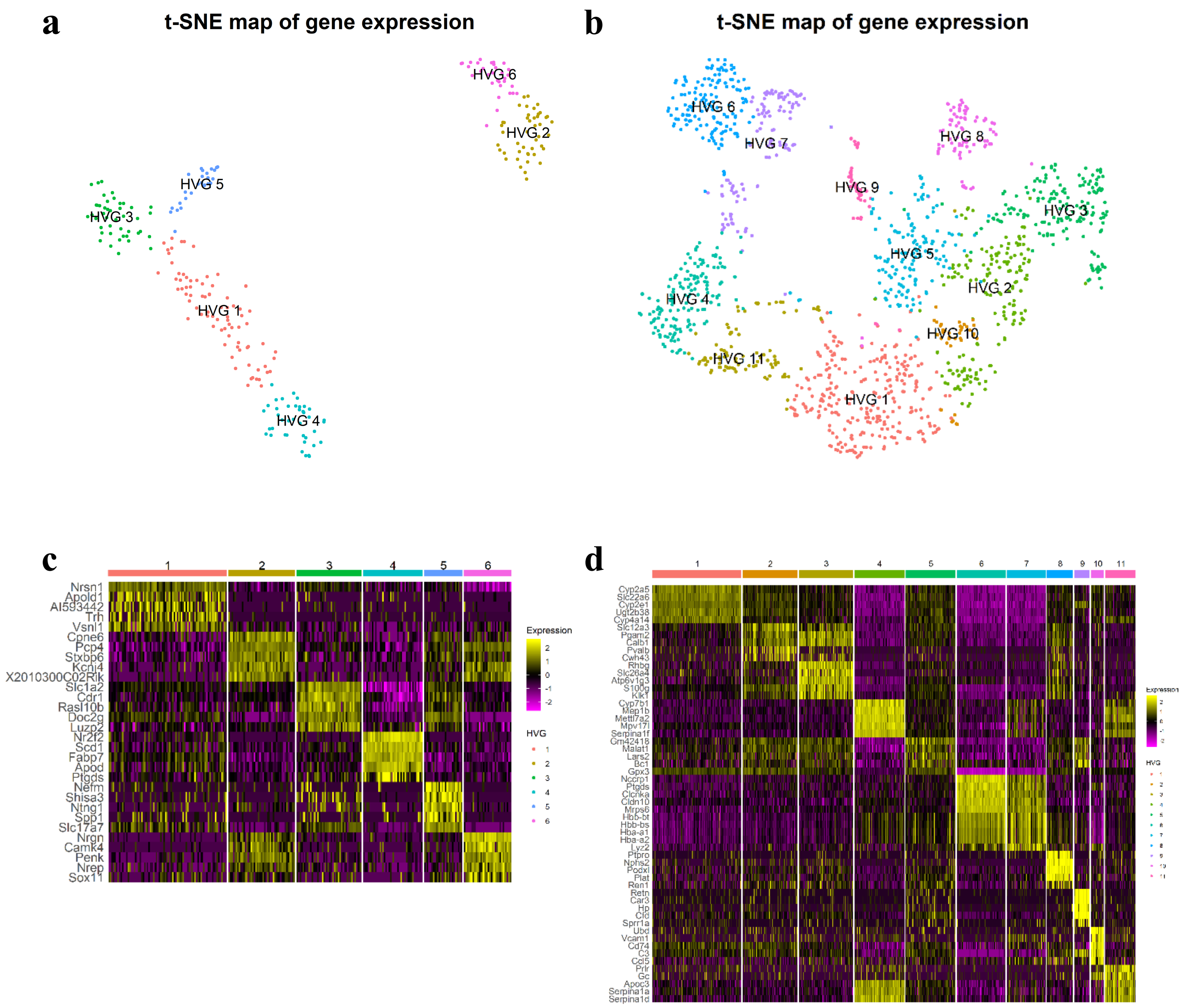


**Supplementary Figure 12. Spot clustering based on HVG in olfactory or kidney tissue.**

(a) t-SNE plot of transcriptomic data from olfactory bulb tissue. HVG-based cluster identity of each spot was visualized. (b) t-SNE plot of transcriptomic data from kidney tissue. HVG-based cluster identity of each spot was visualized. (c) Heatmap showing top 5 differentially expressed genes in each HVG-based cluster from olfactory bulb data. (d) Heatmap showing top 5 differentially expressed genes in each HVG-based cluster from kidney data.

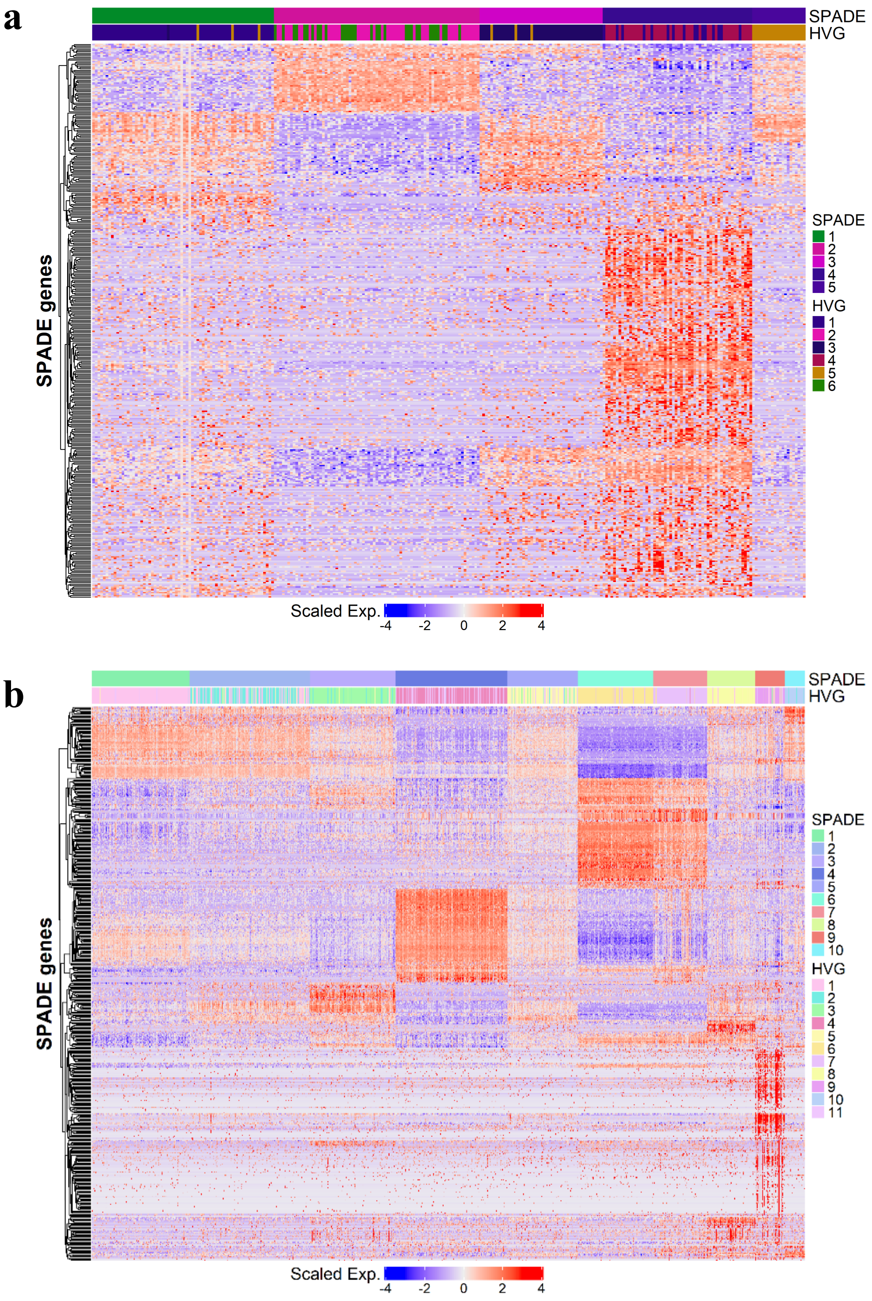


**Supplementary Figure 13. Expression of SPADE genes and clusters of two different datasets.** Heatmaps for expression of SPADE genes in every sampling spot in (a) olfactory bulb or (b) kidney tissue. SPADE or HVG-based cluster identity of each spot was annotated on top of heatmap. Hierarchical clustering for SPADE genes was performed and the results were presented on the left panel.

**a**

**b**

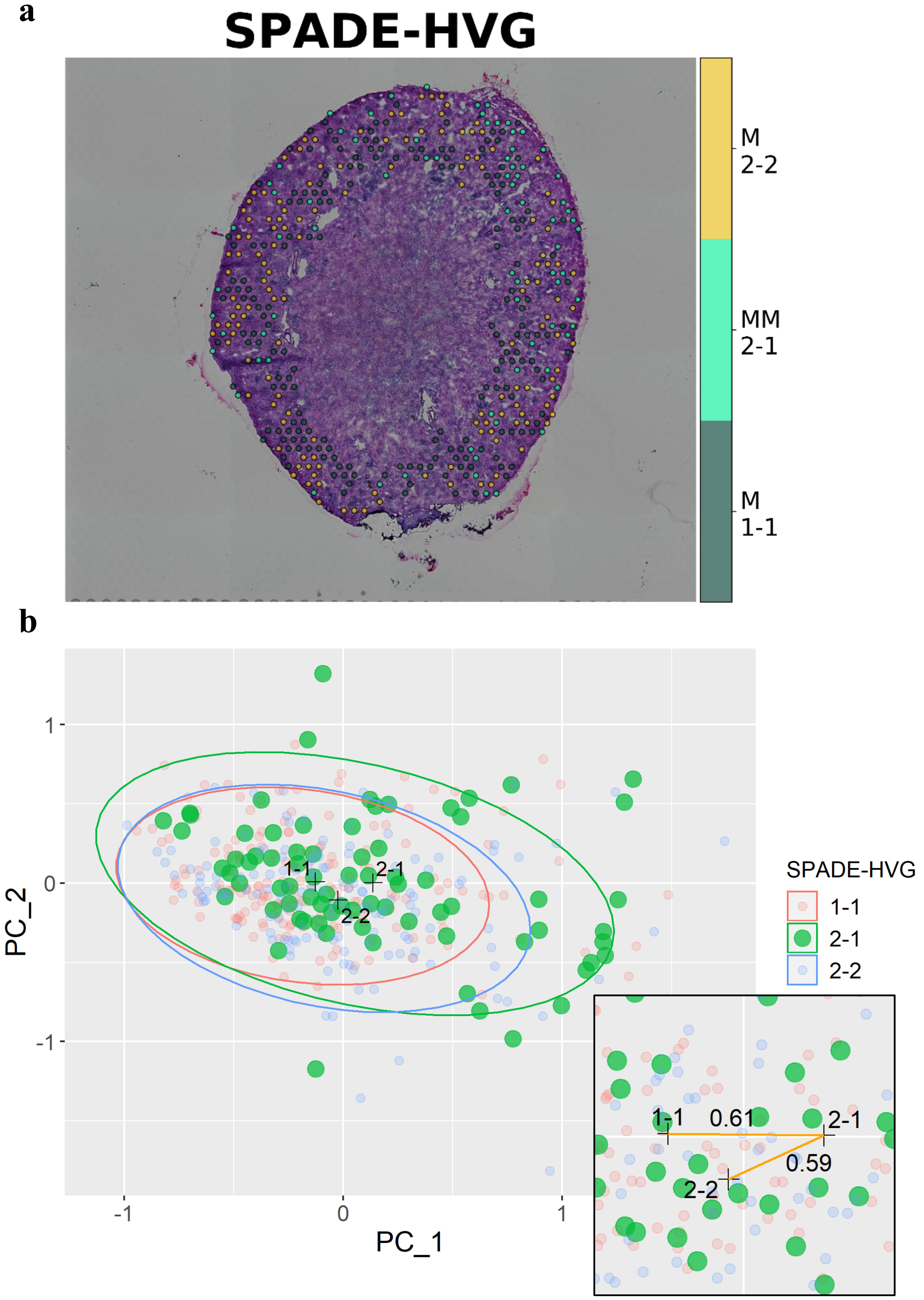
SPADE-HVG
M
2-2
MM
2-1
M
1-1
1
0
-1
-1
0
1
PC_1
PC_2
1-1
2-1
2-2
SPADE-HVG
1-1
2-1
2-2
1-1
0.61
2-1
0.59
2-2

**Supplementary Figure 14. Characteristics of mismatched spot clusters in kidney tissue.**

(a) Spatial mapping of the SPADE 1-HVG 1 and SPADE 2-HVG 2 matched and SPADE 2-HVG 1 mismatched spot clusters in kidney tissue. M is abbreviation for matched cluster and MM is for mismatched cluster. (b) PC1 and PC2 image latent plot for matched and mismatched spot clusters in kidney tissue. A distribution of the mismatched spot clusters were presented in green dots. 95% confidence ellipse for each cluster based on multivariate t-distribution was exhibited on PC plot. Enlarged picture in the bottom shows median distance between mismatched spots and center of mass for matched clusters.
